# Single-Cell RNA-Seq Analysis Reveals Four Stages of Replicative Senescence in Human Umbilical Vein Endothelial Cells

**DOI:** 10.1101/2023.11.12.566783

**Authors:** Minseo Ahn, Junil Kim, Jae Ho Seo

## Abstract

Cellular senescence is a phenomenon marked by an irreversible growth arrest with altered physiological properties. Many studies have focused on the characteristics of cells that have already entered a senescent state. However, to elucidate the mechanisms of cellular aging, it is essential to investigate the gradual transition of proliferative cells into senescent cells. We assumed that cellular senescence is a complex and multi-step process, and each stage exhibits unique traits. To test this hypothesis, we utilized publicly available single-cell RNA-Seq (scRNA-Seq) data from human umbilical vein endothelial cells (HUVECs) undergoing replicative senescence. We employed Seurat and Monocle 3 to capture the transition from proliferating to senescent states in HUVECs. Four clusters were identified, and each cluster displayed distinct expression patterns of cellular senescence markers and the senescence-associated secretory phenotypes (SASPs). We also employed SCENIC to identify the expression patterns of core transcription factors (TFs) during replicative senescence. While the majority of TFs exhibited a linear trend, HMGB1, FOSL1, SMC3, RAD21, SOX4, and XBP1 showed fluctuating expression patterns during replicative senescence. Furthermore, the expression patterns of these TFs exhibited slight differences in the ionizing radiation (IR) model of senescence. Overall, our study unveils the distinct characteristics of each phase during replicative senescence and identifies expression trends in TFs that may play pivotal roles in this process. These findings highlight the intricate nature of cellular senescence and provide new insights into the process of cellular aging.

## 1. Introduction

Cellular senescence represents a condition where cells undergo a permanent cell cycle arrest and develop resistance to apoptosis, and this condition is triggered by diverse stimuli such as DNA damage, telomere shortening, and activation of oncogenes [1,2]. Another characteristic of cellular senescence is that senescent cells secrete senescence-associated secretory phenotypes (SASPs) molecules, which recruit immune cells to clear out senescent cells, induce secondary senescence, and cause chronic inflammation [3]. The accumulation of senescent cells during aging can lead to tissue or organ dysfunction and contribute to various age-related diseases [4]. In 1961, Leonard Hayflick found that when normal human fetal cells are cultured in laboratory conditions after dividing 40–60 times, they eventually become senescent [5]. This phenomenon is commonly referred to as replicative senescence, which is caused by telomere shortening [6]. Replicative senescence, a heterogeneous and multi-stage process, is widely recognized as a key mechanism that significantly contributes to tissue and organ aging.

Most research related to cellular senescence has focused on cells that have already become senescent. However, it is crucial to investigate the transition from a proliferative state to a senescent state to gain a better understanding of cellular aging. Single-cell RNA-Seq (scRNA-seq) technology allows for the investigation of transcriptomes at the single-cell level, making it a powerful tool for studying heterogeneity in cellular dynamics [7,8]. In this study, we utilized scRNA-seq data from human umbilical vein endothelial cells (HUVECs) undergoing replicative senescence to investigate molecular mechanisms in the transition toward replicative senescence. Through scRNA-seq analysis, we identified four clusters that represent transitional states from the proliferative to the senescent state. Our result shows that well-known senescence markers exhibited a monotonic expression pattern, but some SASPs (Senescence-Associated Secretory Phenotype) levels displayed a biphasic pattern with a peak at an intermediate state. We also uncovered transcription factors (TFs) that play a crucial role in replicative senescence using SCENIC [9]. Several TFs were observed to have the highest expression in the intermediate state. Moreover, when comparing the TF patterns with IR-induced senescence, approximately half of the TFs exhibited different expression patterns. These findings suggest that replicative senescence is a complex and multi-stage process.

## 2. Results

### Single-cell RNA-seq analysis reveals four clusters representing the transition from proliferating to senescent states during replicative senescence in HUVECs

To explore the heterogeneity of HUVECs during replicative senescence, we used a scRNA-seq dataset from Zirkel et al. which ∼65% of HUVECs at passage 16 had entered senescence in vitro [10]. The 5,267 cells with at least 1,500 genes detected in each single cell were used for Seurat, which is a standard scRNA-seq analysis platform [11]. Principal component analysis (PCA) and Uniform Manifold Approximation and Projection (UMAP) were conducted for dimension reduction and visualization. We employed unsupervised clustering to group the cells, resulting in the identification of four distinct clusters **(Figure 1A)**. We assumed that some of the cells are in a proliferative state, some are in a senescent state, and the rest are in an intermediate stage transitioning from the proliferative to the senescent state. To test this idea, we identified Differentially Expressed Genes (DEGs) for each cluster **(Figure 1B)**.

**Figure 1.**
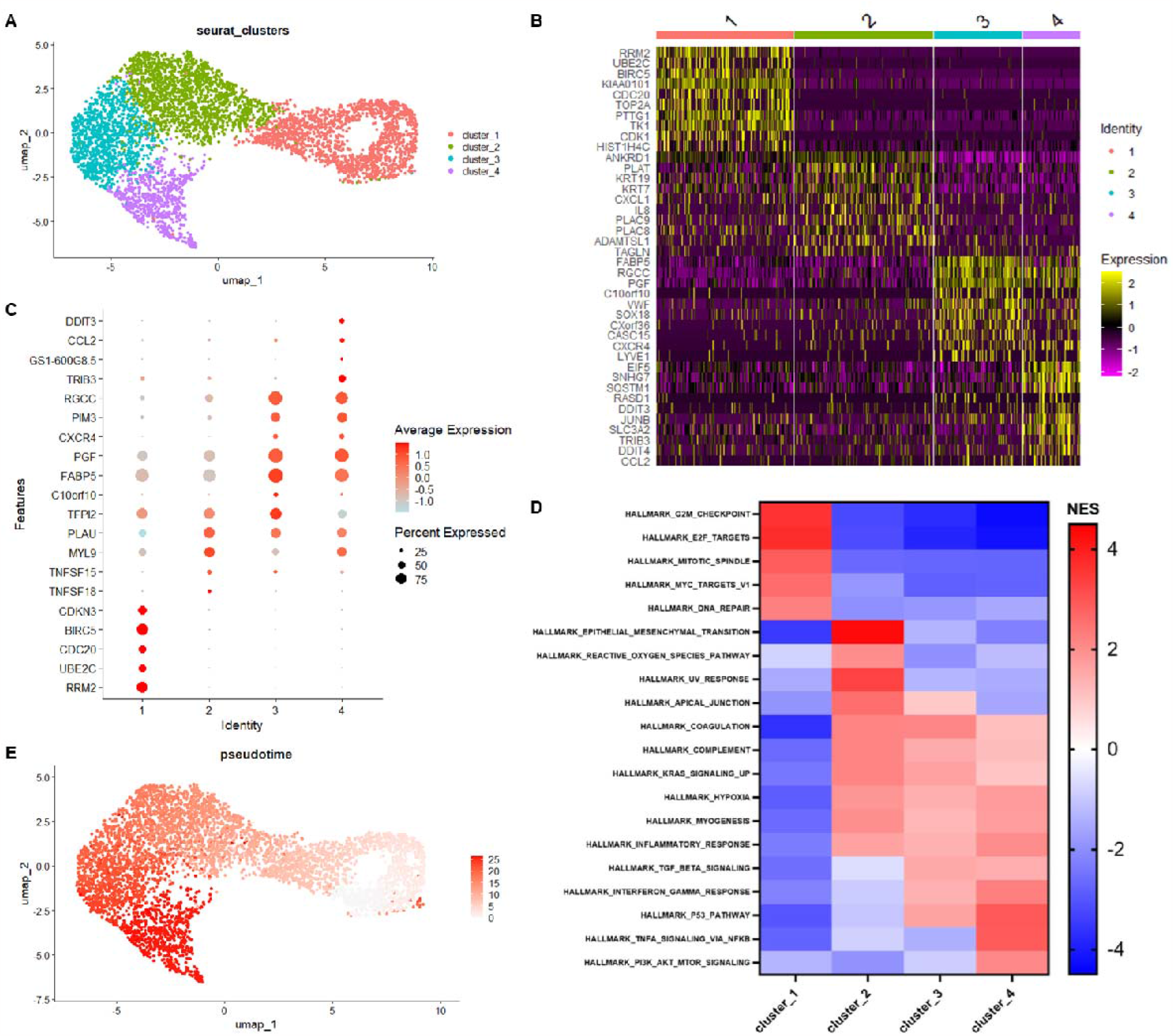
Single-cell RNA-seq analysis reveals four clusters depicting the transition from proliferation to senescence in HUVECs during replicative senescence. **(A)** UMAP plot of HUVECs with four clusters. **(B)** Heatmap of differentially expressed genes of four clusters. Colors close to yellow indicates highly expressed genes while colors close to dark purple represents low expression level. **(C)** Dot plot of top 5 differentially expressed genes (DEGs) for each cluster. **(D)** Heatmap of Gene Set Enrichment Analysis (GSEA) for each cluster. Colors indicate Normalized Enrichment Score (NES) **(E)** UMAP plot of pseudotime value for each cell.

Cluster 1 exhibited elevated expression levels of genes associated with the mitotic cell cycle, such as CDC20 and CDK1 [12,13]. Interestingly, SASP-related genes IL8 and CXCL1 were identified as marker genes for cluster 2. Dot plots with the top 5 variable markers for each cluster effectively distinguished clusters 1 and 2, while clusters 3 and 4 exhibited some overlapped markers **(Figure 1C)**. We also found that cluster 1 exhibited a high enrichment of proliferation-related hallmarks, such as E2F and G2M checkpoints with gene set enrichment analysis (GSEA) [14]. These hallmarks were significantly downregulated starting from cluster 2, indicating cell cycle arrest. In cluster 3, there were high enrichment scores for cellular senescence-related hallmarks, including the p53 pathway and Interferon□gamma (IFN□γ) signaling, and these scores were even higher in cluster 4 [15,16]. Moreover, Cluster 4 exhibited characteristic features of Tumor Necrosis Factor Alpha (TNF-α), which is an inflammatory signaling molecule secreted by cells undergoing senescence [17]. Based on this profiling, we assumed that cluster 1 represents the proliferative state and designated it as the starting point of a senescence trajectory. A trajectory analysis using Monocle 3 highlighted that the pseudotime values were lowest in cluster 1 and highest in cluster 4 (Figure 1E) [18]. Overall, we designated cluster 1 as the proliferative state, cluster 4 as the senescent state, and clusters 2 and 3 as intermediate states for further analysis.

### Four clusters of HUVECs showed distinct senescence markers and SASP profiles

Next, we examined senescence markers within each cluster **(Figure 2A)**. Proliferation markers, such as MKI67 (Ki-67) and PCNA, exhibited high expression in cluster 1 and remained consistently low in other clusters [19,20]. The SA-β-gal activity was indirectly measured by the gene GLB1, which encodes lysosomal β-D-galactosidase [21,22]. In our data, GLB1 levels gradually increased during replicative senescence. Reduced expression of the nuclear lamina protein Lamin B1 (LMNB1) was also observed during replicative senescence [23]. Subsequently, we examined genes associated with the p53 pathway, such as p53, p21, and p16, which are well-known regulators of cellular senescence [24]. As expected, their expression levels increased from cluster 1 to cluster 4. We then assessed the levels of ATM and ATR, which are involved in the DNA damage response [25]. Unexpectedly, ATM levels continued to rise until cluster 3 and decreased in cluster 4, while ATR levels were highest in cluster 1, decreased until cluster 3, and then increased again in cluster 4.

**Figure 2.**
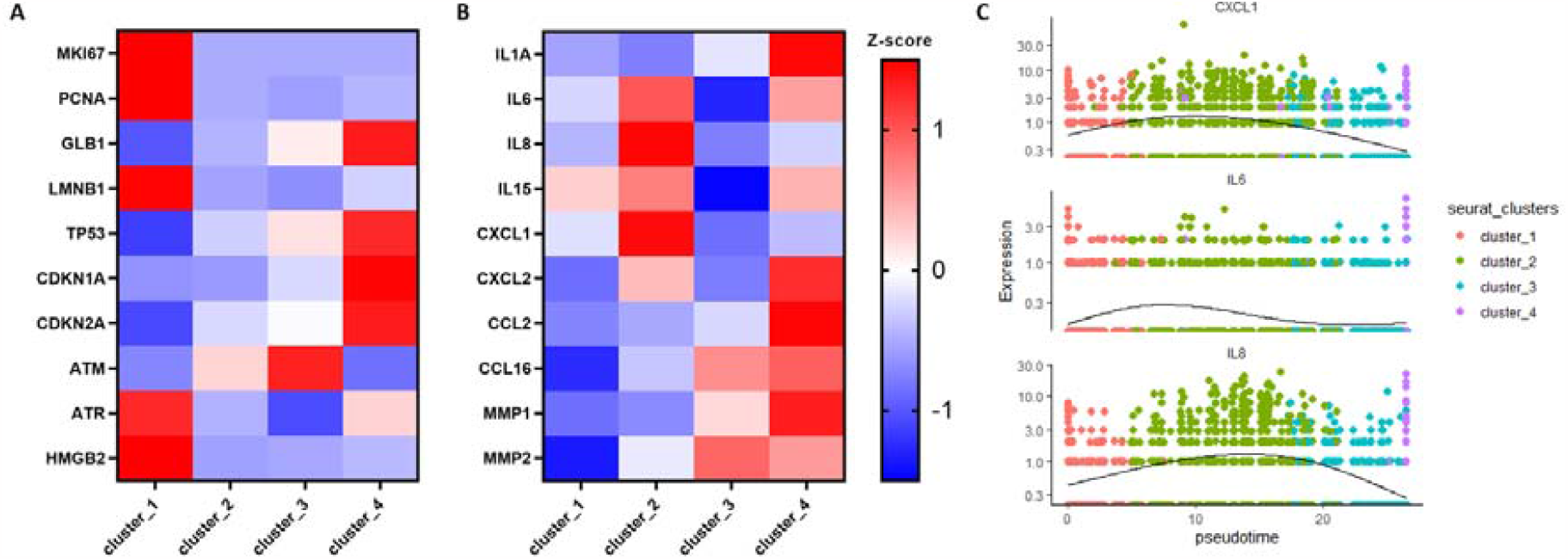
Four HUVEC clusters exhibited unique senescence markers and SASP marker profiles. **(A, B)** Heatmap of senescence markers and senescence-associated secretory phenotypes (SASPs) markers. **(C)** Expression patterns of SASP markers that showed biphasic patterns during replicative senescence. The X-axis corresponds to the pseudotime value while the Y-axis corresponds to the expression level.

Chromatin in aged cells undergoes a reorganization process to create structures known as senescence-associated heterochromatin foci (SAHF) [26]. Zirkel et al. found that HMGB2 plays a crucial role in regulating the chromatin structure of SASP gene loci [10]. The level of HMGB2 decreased as replicative senescence progressed in our data. These results further support that the clusters and pseudotime analysis effectively capture the dynamics of replicative senescence. Senescent cells secrete SASP markers, which include various inflammatory molecules and growth factors [3]. SASP factors influence neighboring cells frequently associated with inflammatory responses and contribute to secondary senescence. We examined the expression level of SASP factors, including cytokine, chemokine, and extracellular protease in each cluster. Interestingly, while about half of the SASP genes exhibited higher expression levels as they progressed toward cluster 4, CXCL1, IL6, and IL8 showed their highest expression in cluster 2. Trajectory analysis also displayed that the expression levels of these genes were notably elevated in the intermediate stage of the pseudotime, which corresponds to cluster 2. This suggests that some SASP components may be produced more abundantly during intermediate phases of replicative senescence.

### Regulatory network analysis uncovers distinct expression patterns of TFs during HUVECs replicative senescence

Since cell fate is greatly influenced by TFs, we used SCENIC to construct a regulatory network of HUVECs during replicative senescence and identify core TFs [27]. By employing the SCENIC, we identified the top 25 variable TFs **(Figure 3A)**. Some of the TFs identified through SCENIC have been known to be associated with cellular aging in earlier studies. For example, HMGA1 and HMGA2 are reported to specifically accumulate on senescent cell chromatin [28]. Also, many studies have highlighted the association of SOX TFs with aging and age-related diseases [29]. In addition, sustained activation of JunB in fibroblasts disrupts interactions within the stem cell niche, leading to the promotion of skin aging [30]. These findings support the idea that identified TFs have a significant link to cellular senescence.

**Figure 3.**
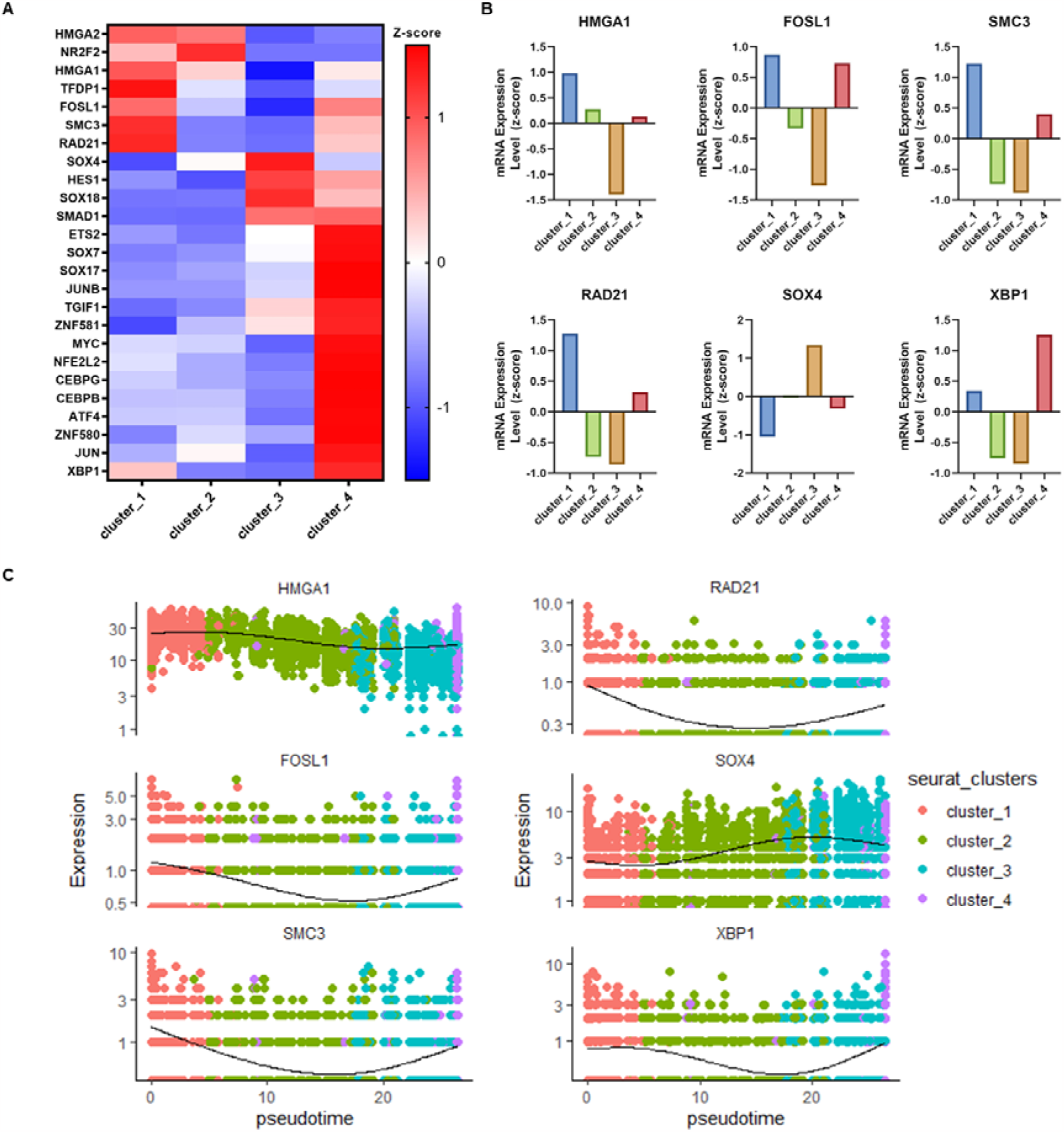
Regulatory network analysis reveals distinct transcription factor expression patterns during HUVEC replicative senescence. **(A)** The top 25 variable transcription factors (TFs) identified by pySCENIC. **(B)** Bar plots of TFs that showed biphasic pattern during HUVECs replicative senescence. **(C)** Expression patterns along with pseudotime values of 6 TFs showed in Figure 3B.

Most TFs showed a monotonic expression pattern, with their levels either increasing or decreasing as the cells entered senescent. Additionally, the majority of these TFs displayed high expression in cluster 4, which is closely associated with the senescent state. On the other hand, FOSL1, HMGA1, SMC3, RAD21, SOX4, and XBP1 exhibited biphasic patterns in their expression levels **(Figure 3B)**. For example, in the case of FOSL1, the expression level continued to decrease until cluster 3 and then increased again in cluster 4. Similarly, for SOX4, the expression level increased as it transitioned from cluster 1 to 3, and then decreased from 3 to 4. The biphasic expression patterns of these TFs also appear at the single-cell level trajectory **(Figure 3C)**, indicating that replicative senescence consists of multiple stages, each of which may exhibit distinct characteristics.

Notably, previous studies have demonstrated that HMGA1 cooperates with p16 to promote SAHF formation and trigger cell cycle arrest in IMR90 fibroblasts undergoing oncogene-induced senescence [31]. Our data reveals a fluctuating pattern in HMGA1 levels, initially peaking in cluster 1, decreasing until cluster 3, and increasing in cluster 4. This suggests the possibility that a high level of HMGA1 may have different functions in proliferative cells. Furthermore, cohesin plays a crucial role in establishing the connection between duplicated sister chromatids, making it a vital component for the proper separation of chromosomes during cell division [32]. It has been reported that there is a notable increase in the expression of genes that are enriched in the subtelomeric regions during replicative senescence in human lung primary fibroblasts [33]. This includes genes like RAD21 and SMC3. Interestingly, the expression levels of RAD21 and SMC3 were high in cluster 1, decreased in the intermediate state, and increased again as cells progressed towards the senescent state. This suggests the potential significance of cohesin activity in the replicative senescence.

### The expression patterns of TFs associated with IR-induced cellular senescence differ from those observed in replicative senescence in HUVECs

Senescent cells exhibit distinct profiles depending on the induction method. To compare the expression patterns of TFs associated with replicative senescence to those induced by other methods, we utilized RNA-Seq data from GSE130727, where cellular senescence was induced in HUVECs using ionizing radiation (IR) [34]. In this dataset, we computed the fold change for the previously identified top 25 TFs between the control group and the group exposed to ionizing radiation (IR). Likewise, we calculated the fold change for the top 25 TFs between cluster 1, representing a state close to the proliferative stage, and cluster 4, representing a state close to the senescent stage, and compared these fold changes. During cellular senescence induced by two different methods, 13 out of 25 TFs (52%) exhibited a consistent trend, while 12 out of 25 TFs (48%) displayed a different pattern **(Figure 4)**. This suggests that cellular senescence occurs through different pathways depending on the method of senescence induction.

**Figure 4.**
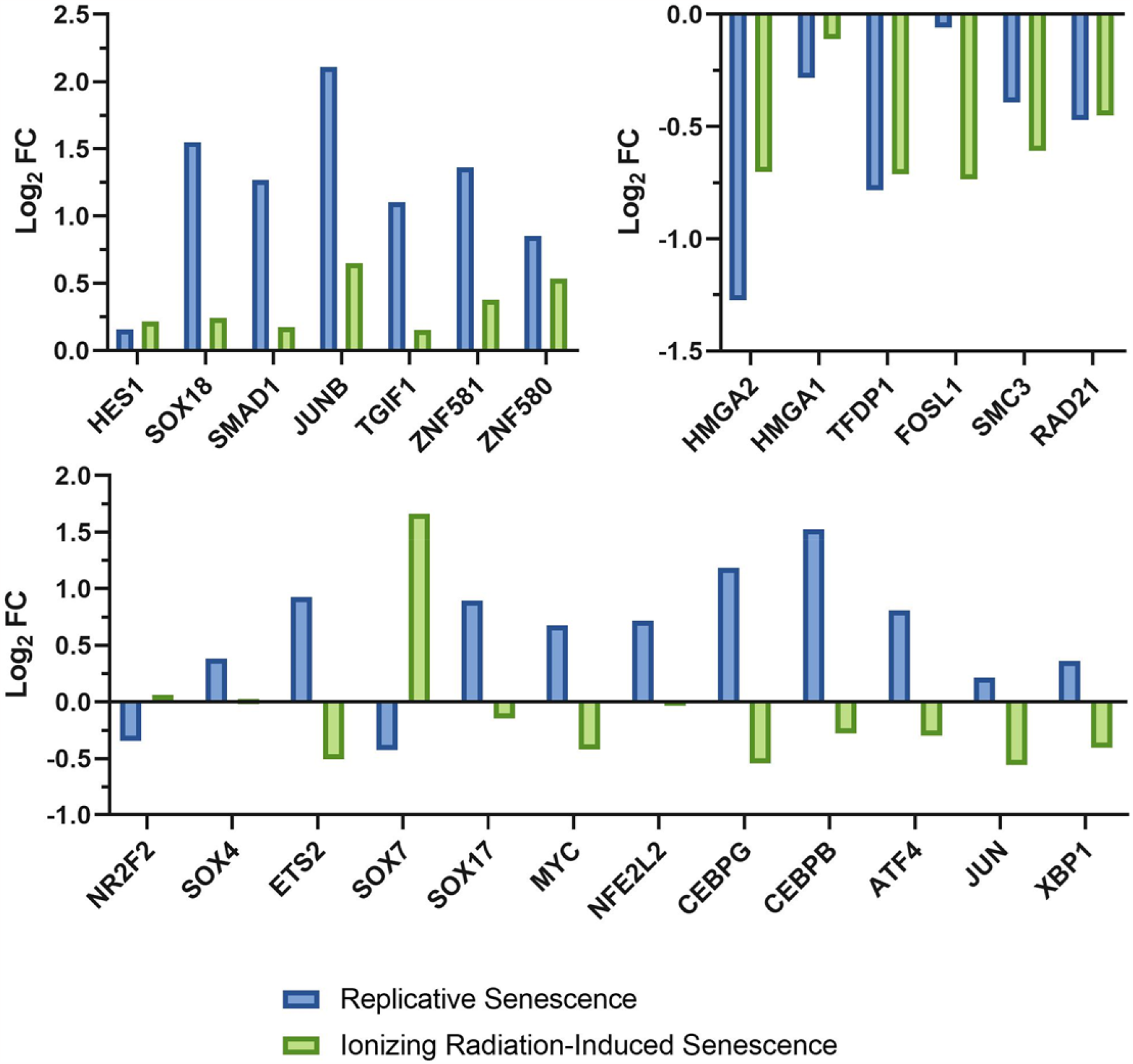
Profile of transcription factors differ in ionizing radiation-induced senescence compared to replicative senescence in HUVECs. Fold changes for the top 25 variable transcription factors (TFs) in replicative senescence were calculated by comparing expression levels of TFs in cluster 1 and cluster 4. Fold changes of these TFs in ionizing radiation-induced senescence were computed by comparing the control group and the IR-exposed group.

Many studies demonstrated that members of the SOX gene family showed changed expression patterns as cells and tissues age [29]. Interestingly, among the analyzed SOX family transcription factors, except for SOX18, there were distinct expression patterns observed in replicative senescence and IR-induced senescence. These results imply that SOX family genes may be regulated differently depending on the type of cellular senescence. Furthermore, CCAAT-enhancer-binding proteins (C/EBPs) are transcription factors that promote chromatin modification and recruit basal TFs with co-activators like CBP. Intriguingly, C/EBPβ (CEBPB) and C/EBPγ (CEBPG), which belong to the C/EBPs family, exhibited a significant increase during replicative senescence, whereas minimal to no significant changes were observed in the IR-induced senescence model [35]. This indicates that the C/EBPs family may play an important role in replicative senescence but not in IR-induced senescence.

## 3. Discussion

Cellular senescence involves a permanent state of cell cycle arrest. In somatic tissues, telomerase expression decreases at the end of embryogenesis, resulting in telomere shortening with each cell division and ultimately leading to replicative senescence. However, the transition process from a proliferative state to a senescent state has been not fully studied. To gain a deeper understanding of the aging process, it is essential to investigate the events occurring during replicative senescence. In this study, we utilized scRNA-seq analysis to investigate the transition from proliferative to senescent states in HUVECs during replicative senescence. Unsupervised clustering identified four distinct clusters, and subsequently, we conducted DEG and GSEA analysis to elucidate the properties of each cluster. Cluster 1 showed elevated expression of genes related to the mitotic cell cycle, indicating a proliferative state. GSEA revealed that cluster 1 was enriched in proliferation-related hallmarks, while these hallmarks were significantly downregulated from cluster 2 onwards, indicating cell cycle arrest. Clusters 3 and 4 exhibited high enrichment scores for cellular senescence-related hallmarks, such as the p53 pathway, TNF-α, and IFN-γ signaling. Based on these results, cluster 1 was designated as the proliferative state, cluster 4 as the senescent state, and clusters 2 and 3 as intermediate states.

Next, we investigated individual markers of senescence SASP for each cluster. Interestingly, ATM, which is one of the markers of DNA damage response, and some SASP markers including CXCL1, IL6, and IL8 are highest in intermediate states. We also performed regulatory network analysis using SCENIC and identified core TFs that may have a critical role in replicative senescence. Remarkably, the expression levels of FOSL1, HMGA1, SMC3, RAD21, SOX4, and XBP1 displayed varying patterns throughout the process of replicative senescence. This suggests that expression levels of some senescence markers, SASP markers, and transcription factors do not simply increase or decrease as cellular senescence progresses, but they exhibit various patterns in the intermediate state. This implies that replicative senescence is complex and involves multiple processes.

We also identified several TF families that play roles in multiple biological processes or pathways. RAD21 and SMC3, associated with cohesin, exhibited decreased expression in the intermediate state but increased expression upon transitioning to the senescent state. Certain genes from the SOX and C/EBP family were recognized as key TFs during replicative senescence in HUVECs and their expression patterns differed between replicative senescence and ionizing radiation-induced senescence. Our study provides insight into the intricate and multifaceted nature of cellular senescence. This study suggests that the production of different types of SASP markers can vary depending on the cellular state during cellular senescence. Additionally, it provides a list of several TFs that may play a crucial role in replicative senescence.

However, further studies are needed to confirm whether these TFs play crucial roles in other cellular senescence induction methods, such as oncogene-induced senescence, or in different cell lines with biological experiments.

## 4. Methods

### Single-cell RNA-seq Data Analysis

The scRNA-seq data of HUVECs at passage 16 was obtained from the NCBI Gene Expression Omnibus (GEO) repository under the accession number GEO: GSE98448. R (v.4.2.3) and RStudio (2023.06.1+524) were used for most analyses. The 5,267 cells with at least 1,500 genes were used for the scRNA-seq analysis pipeline of Seurat (v4.9.9.9060). Cells that have unique feature counts over 5000 or less than 1500 and have more than 6% mitochondrial counts were filtered out. The data was normalized and 5,000 variable genes were identified with Seurat default setting.

### Dimension Reduction and Clustering Analysis

Principal component analysis (PCA) and Uniform Manifold Approximation and Projection (UMAP) were performed for dimension reduction and visualization. The first 10 PCs were used for UMAP and clustering was determined by the elbow method. Cells were clustered using a network-based clustering method implemented in Seurat.

### DEGs and Gene Set Enrichment Analysis

DEGs of each cluster were identified using the *FindAllMarkers* function implemented in Seurat with p<0.01. The heatmap of the top 10 DEGs was produced with the *DoHeatmap* function in Seurat. Genes were ranked with average log 2 foldchange values and Gene set enrichment analysis (GSEA) was performed with the fGSEA package. Hallmarks of the gene set from MSigDB were used for the analysis.

### Trajectory Analysis

Monocle 3 (v.1.3.1) was used for trajectory analysis. The Seurat object of HUVECs data was converted to a CellDataSet object with SeuratWrappers (v.0.3.1). The CellDataSet object was pre-processed and dimension reduction was performed with Monocle 3 default setting. Single-cell trajectories were constructed with *learn_graph* and *order_cells* function in Monocle 3. Pseudotime values were extracted from the CellDataSet object and applied to the Seurat object to draw plots.

### Regulatory network analysis with SCENIC

Python (v.3.10.9) and pySCENIC (v.0.12.1) were used for regulatory network analysis. 5000 variable genes of HUVECs during replicative senescence were used for the analysis. The TFs were identified with pySCENIC default settings. 156 TFs were identified with pySCENIC and the top 25 variable TFs were used for further analysis.

### Ionizing radiation-induced senescence RNA-Seq data analysis

RNA-Seq data from HUVECs that are exposed to 4 Gy of ionizing radiation was obtained from the repository under the accession number GEO: GSE130727. The fold change of core TFs in IR-induced senescent cells was determined by comparing the average expression levels of these TFs for three samples in both the control group and the IR-exposed group.

